# Overexpression of *SHORT-ROOT2* transcription factor enhanced the outgrowth of mature axillary buds in poplar trees

**DOI:** 10.1101/2021.08.08.455544

**Authors:** Minglei Yi, Heyu Yang, Shaohui Yang, Jiehua Wang

## Abstract

Plant branching is usually prevented by an actively proliferating apex. In poplars, one GRAS family member, *SHORT-ROOT2* (*PtSHR2*), was preferentially expressed in axillary buds (AXBs) and was inducible during bud maturation and activation. Overexpression of *PtSHR2* (*PtSHR2*^OE^) in hybrid poplar impaired the apical dominance and simultaneously promoted the outgrowth of axillary branches below the maturation point (BMP), accompanied by regulated expression of genes critical for axillary meristem initiation and bud formation.
Following a detained phenotypic characterization, we compared the IAA and *trans*-zeatin levels in apical shoots and AXBs of wild-type and *PtSHR2*^OE^ trees, together with gene expression analyses and defoliation, decapitation, and hormone reapplication assays.
*PtSHR2*^OE^ AXBs contained a significantly lower ratio of auxin to cytokinin than wild-type AXBs, particularly in those below the BMP. Decapitation induced a faster bud burst in *PtSHR2*^OE^ trees than in wild-type plants, and it could be strongly inhibited by exogenously applied auxin and cytokinin biosynthesis inhibitor, but only partially inhibited by N-1-naphthylphthalamic acid (NPA).
An impaired basipetal auxin transport, rather than an insufficient auxin biosynthesis or auxin insensitivity, disturbed the local hormonal homeostasis in *PtSHR2*^OE^ AXBs, which in turn enhanced the axillary bud initiation and promoted the bud release.

**Highlight:** Overexpression of *PtSHR2* in poplar impaired the apical dominance and promoted axillary bud outgrowth below the maturation point through disturbing the basipetal auxin transport and auxin and cytokinin homeostasis.

## Introduction

In seed plants, when the shoot apex is proliferating actively during the growth season, axillary meristems (AMs) usually develop into quiescent axillary buds (AXBs) in the leaf axil due to the apical dominance (Cline, 1991, 1997; Leyser, 2005; Phillips, 1975; Thimann and Skoog, 1934). In deciduous woody perennials, the development of AXBs is completed at the bud maturation point (BMP), at which position they contain a predictable number of embryonic leaves. Then, the buds undergo different dormant states throughout the year in response to environmental changes and are susceptible to grow in response to mild temperatures of the following year (Rohde and Bhalerao, 2007).

For plants under active growth, the paradormancy of AXBs is achieved at least in part through the local biosynthesis of auxin and its basipetal transport (Bennett and Leyser, 2006), which has generated two non-exclusive models. In the auxin canalization model, auxin in the main stem competitively inhibits an auxin export flux from AXBs towards stems, which is a prerequisite for axillary bud outgrowth (Bennett *et al*., 2014; Sachs, 1981). In the other model, cytokinin and strigolactones, two phytohormones inducing and inhibiting bud outgrowth, respectively; are proposed as second messengers for the apically-derived auxin to indirectly inhibit bud outgrowth (Aguilar-Martínez *et al*., 2007; Braun *et al*., 2012; Dun *et al*., 2012; Gomezroldan *et al*., 2008; Hayward *et al*., 2009; Snow, 1937; Tanaka *et al*., 2006; Umehara *et al*., 2008). When the dominant auxin source is removed by decapitation, the local synthesis of cytokinin in buds is stimulated, which upregulates the polarization of PIN proteins and facilitates the auxin export (Balla *et al*., 2011; Domagalska and Leyser, 2011; Kalousek *et al*., 2010). Notably, the induced outgrowth of buds can be prevented again by applying auxin to the decapitation site (Thimann and Skoog, 1933).

With the name representing three family members including GAI (GIBBERELLIC ACID INSENSITIVE), RGA (REPRESSOR of GAI) and SCR (SCARECROW), GRAS-domain transcription factors represent a plant-specific transcription regulator family playing important roles in multiple aspects of plant organogenesis. However, the involvements of GRAS proteins in branching process have not been revealed extensively, and only one Arabidopsis GRAS protein, LATERAL SUPPRESSOR (LAS), has been shown to be required to establish AMs since the *las* mutant fails to initiate AM during vegetative development (Greb *et al*., 2003). As the most prominent member of the GRAS family, SHORT-ROOT (SHR) has been most intensively studied in *Arabidopsis thaliana* root, where it is believed to physically interact with another related GRAS protein, SCARECROW (SCR), to regulate stem cell maintenance and radial patterning (Di Laurenzio *et al*., 1996; Helariutta *et al*., 2000; Sabatini *et al*., 2003). In both *shr* and *scr* mutants, the quiescent center (QC) is disorganized and proliferating cells are prematurely depleted, resulting in only a single layer of ground tissue (Di Laurenzio *et al*., 1996; Helariutta *et al*., 2000). Notably, the regulation of tissue patterning by SHR in Arabidopsis involves the modulation of phytohormone homeostasis and higher endogenous levels of both cytokinin and auxin were observed in roots of *shr* mutant than those in the wild type (Lucas *et al*., 2011; Moubayidin *et al*., 2013). In addition to root, SHR and SCR also cooperate to regulate cell proliferation and differentiation in shoot organs (Cui *et al*., 2014; Dhondt *et al*., 2010; Gao *et al*., 2014; Miguel *et al*., 2015; Slewinski *et al*., 2012). Loss of SHR function in Arabidopsis causes retarded leaf growth and stunted shoot phenotype, which is believed to be independent from the impaired root development (Dhondt *et al*., 2010). Despite of the above-mentioned results, roles of SHR involved in AM establishment or AXB development and activation has not been reported so far.

In Populus genome, three *SHR* genes, namely *PtSHR1* and *PtSHR2A*/*2B*, have been previously identified (Wang *et al*., 2011). *PtSHR1* shares the highest similarity with *AtSHR* and is specially expressed in the vascular cambial zone (Schrader *et al*., 2004; Wang *et al*., 2011) where it regulates cambial cell division and lateral meristem activity (Koizumi *et al*., 2012; Wang *et al*., 2011). Partial suppression of *PtSHR1* in transgenic poplars led to taller trees with a larger vascular cambial zone due to an increased meristem cell proliferation (Wang *et al*., 2011). With shorter 5’ ends when compared with *AtSHR*, *PtSHR2A* and *PtSHR2B* encode almost identical proteins and PtSHR2B has been reported to function in the phellogen and regulate the phellem and periderm formation during secondary growth, which possibly act through modulation of cytokinin homeostasis (Miguel *et al*., 2015).

In this work, we overexpressed *PtSHR2B* in hybrid aspen clone INRA 717-1B4 and found that in addition to their stunted primary growth, *PtSHR2B* overexpression *(PtSHR2B*^OE^) trees exhibited another remarkable phenotypic character, e.g., an enhanced AXB outgrowth and high sylleptic branching development with an unambiguous starting point at the BMP. Such release of axillary buds was synchronized with the transition of a leading shoot tips into a terminal bud (TB). Meanwhile, the antagonizing action of auxin and cytokinin in *PtSHR2B*^OE^ trees were dramatically regulated at shoot apex and in axillary buds, which were accompanied with distinctly regulated expression of a set of transcriptional factors controlling AM formation and AXB outgrowth. By performing defoliation, decapitation, and auxin reapplication assays on transgenic and wild-type trees, we demonstrated that the mechanism underlying the bud-flush of *PtSHR2B*^OE^ trees probably included a disrupted basipetal transport of auxin, an enhanced local biosynthesis of cytokinin and a boosted auxin efflux from AXBs. Overall, results presented here not only revealed a novel role of SHR in regulating branching in woody perennials through a highly conserved mechanism as in annual Arabidopsis bud outgrowth, but also provided new evidence for the previous contention that bud development at terminal and axillary positions follows a shared developmental pattern (Rinne *et al*., 2015) and under certain conditions, their identities could be exchanged by disturbed hormonal homeostasis.

## Materials and Methods

### Plant materials and growth conditions

The hybrid poplar (*P. tremula* × *P. alba* clone INRA 717-1B4) was propagated and maintained by transferring shoot segments of 3-4 cm with 1-2 young leaves to fresh one-half Murashige and Skoog (Beijing, China, Sigma-Aldrich). Plantlets were grown at 23 ℃ under a 16 h light/8 h dark cycle with a light intensity of illumination of 5000 lux provided by cool white fluorescent lamp tubes. Then, one-month-old plantlets were transplanted into the nutrient soil and kept in the growth chamber of 70% humidity under 16 h light/8 h dark photoperiod at 25 °C for a minimum of 3 months prior to analysis. For gene expression analysis, apices, axillary buds at various nodes of greenhouse poplars were carefully collected, frozen in liquid nitrogen and store at −80 ℃ prior to RNA extraction. For gene expression analysis during bud activation, decapitation was performed either at the shoot apex or at the BMP site, and the nearest buds were collected from several trees at day 0, 3, 7 and 9.

### Vector construction and poplar transformation

The full-length cDNA of *PtSHR2B* was amplified by PCR using the genomic DNA of *P. trichocarpa*, inserted into *CaMV* 35S promoter and further recombined into the binary vector pBI121 to produce the *35S::PtSHR2B* constructs. Based on the previously reported method (Nilsson *et al*., 1992), all vector constructs were transferred into *Agrobacterium tumefaciens* C58 strain pMP90 and then used to transform the hybrid poplars (*P. tremula × P. alba* clone INRA 717-1B4). In order to construct *PtSHR1pro: GUS* and *PtSHR2Bpro:GUS* transgenic poplars lines, the promoter regions of *PtSHR1* and *PtSHR2B* genes were amplified from the genomic DNA of *P. trichocarpa* and were fused with *uidA* (β-glucoronidase/GUS) reporter gene, respectively; before being transformed into the host plants. The positive transformants were selected on kanamycin-containing media and identified by PCR genotyping with the primers of *CaMV* 35S promoter.

### Sample preparation and histological assay

GUS activity of *SHRpro::GUS* transformants was assayed in paraffin-embedded thin sections according to methods previously reported (Johansson *et al*., 2003). Briefly, transverse hand-cut shoot apices about 2 mm thick were prefixed in cold 90% acetone and then incubated in 1 mM X-glucuronide (X-Gluc) in 50 mM sodium phosphate pH 7.0 and 0.1% Triton X-100, 1 mM K_3_Fe(CN)_6_ and 1 mM K_4_Fe(CN)_6_. They were then fixed in FAA, dehydrated in ethanol, incubated 2 h each in a series of ethanol/xylene mixtures and finally in 100% xylene. After this, samples were infiltrated in 1.5 volumes of xylene and one volume of melted paraffin at 37 °C for about 42 h, and then embedded in paraffin at 62 °C. The embedded samples were sectioned transversely with a rotary microtome at 10 μm before that the cellular location and intensity of the GUS staining was assessed with a light microscope (Zeiss, Shanghai, China). For structural and histological analysis of AXBs, AXBs and adjacent primary stem segments from 3-month-old *PtSHR2*^OE^ trees and WT were fixed in FAA solution, gradually dehydrated, embedded in 0.6-0.7 % (w/v) agarose, sliced with the thickness as 50 μm by a vibratome (Leica VT1000 S, Shanghai, China) and stained with phloroglucin-HCl staining as described before (Sakurai *et al*., 2001).

### Gene expression analyses by qRT-PCR

Total RNA was extracted using the EasyPure Plant RNA Kit (TransGen Biotech, Beijing, China) and 1 μg sample of total RNA was used to synthesize the cDNA using the EasyScript First-Strand cDNA Synthesis SuperMix (TransGen Biotech). The quantitative RT-RCR analysis were performed using SYMR Green fluorochrome technique on a PikoReal 96 Real-time Thermal Cycler (Thermo Fisher Scientific, Shanghai, China) under the following conditions: initial incubation at 95 ℃ for 2 min, followed by 45 cycles of 20 s at 95 ℃, 20 s at 58 ℃ and 30 s at 72 ℃. All primer sequences used were listed in Table S1 and *PtaActin* was used as the internal reference gene and the 2^(−ΔΔCt)^ method was used to analyze the data.

### Quantification of IAA and zeatin content

Zeatin and free IAA contents were measured according to a previously described method (Vitti *et al*., 2013). An explant (300 mg) of shoot or root tissue was ground into powder with liquid nitrogen and add 0.5 mL extraction solvent (2-propanol/H_2_O/formic acid 2: 1: 0.002 in volume) was added. The tubes were sonicated for 30 min at 4 °C. To each tube, 1 mL of dichloromethane (chloroform) was added, and then the samples were shaken for 30 min at 4 °C and centrifuged at 13,000× g for 5 min. After centrifugation, two phases were formed, with plant debris between the two layers, so 1.0 mL of the solvent from the lower phase was transferred using a Pasteur pipette into a screw-cap vial, and the solvent mixture was concentrated using an evaporator with nitrogen flow. Finally, the samples were re-dissolved in 0.2 mL 5% methanol and stored at −20 °C before quantitative analysis. Chromatographic separation was accomplished using a Waters ACQUITY UPLC HSS T3 (100 mm × 2.1 mm, 1.8 um) with the mobile phase of 0.1% formic acid in water (Solvent A) and 0.1% formic acid in methanol (Solvent B) delivered in gradient elution mode at a flow rate of 0.3 mL min^−1^. The elution program used was as follows: 0–1 min, 30% B; 1–5 min, 80% B; 5–10 min, 100% B; 10–15 min, 30% B. The LC eluent was introduced into an AB Sciex 6500+ quadrupole ion trap mass spectrometer (QTRAPMS) (AB Sciex, Framingham, MA, USA) for MRM with electrospray ionization (ESI). Spectra of IAA and *t*-ZR were acquired in the positive ionization mode. Final MS and MRM parameters were as follows: Curtain gas (30 psi), collision gas (medium), IonSpray voltage (5500 V), source temperature (450 °C), ion source gas 1 (50 psi), ion source gas 2 (50 psi). Quantification was performed by MRM of the protonated precursor molecular ions [M + H]^+^ and the related product ions. Chromatograms and mass spectral data were acquired and processed using SCIEX OS-MQ software (AB Sciex, Framingham, MA, USA). Pure standards of each phytohormone (Duchefa Biochemie, Netherlands) were used for identification analysis.

### IAA, NPA and Lovastatin application experiments

For exogenous IAA application, lanolin containing 6 mg g^-1^ NAA (final ethanol concentration 10 %) was applied to the decapitated stump of wild-type and transgenic trees at three sites (below apex, at BMP and at the basal node). As controls, two types of trees were simultaneously treated with only ethanol in the lanolin but lacking NAA. For treatment of intact trees, 30 µM N-1-naphthylphthalamic acid (NPA) dissolved in DMSO (Bae *et al*., 2020) or 5 mM Lovastatin, an inhibitor of cytokinin synthesis, dissolved in ethanolic NaOH [15 % (v/v) ethanol, 0.25 % (w/v) NaOH] (Crowell and Salaz, 1992) was directly applied onto the axillary buds. For treatment of trees decapitated at BMP sites, a 10 mm wide skin ring was ground 1cm below the cut site to remove the wax layer. The worn area is covered with 30 μM NPA in lanolin and then wrapped with parafilm and light-blocking foil (Wan *et al*., 2006). NPA in lanolin and Lovastatin were applied directly over the decapitation site, and then covered as above-mentioned. Control plants were similarly treated without NPA or Lovastatin.

## Results

### AXB-specific expression and decapitation induction of *PtSHR2B* in hybrid poplar

In the genome of *Populus trichocarpa*, there are three AtSHR homologs, namely PtSHR1 and PtSHR2A/2B. Due to their 94% nucleic acid sequence identity and 96% amino acid similarity, PtSHR2A and PtSHR2B are collectively denoted as SHR2. When the mRNA levels of *PtaSHR* genes were analyzed by qRT-PCR in different tissues of hybrid poplar (*Populus tremula* × *P. alba*) clone INRA 717-1B4, both *PtaSHR1* and *PtaSHR2* were transcribed at high levels in apical shoots and roots; and in petioles and stems, *PtaSHR2* exhibited higher transcriptional abundances than *PtaSHR1* (Fig. 1A). In shoot tips of transgenic hybrid poplar expressing a transcriptional fusion between upstream sequence of *PtSHRs* and the *uidA* (ß-glucoronidase/GUS) reporter gene, histochemical GUS activity was located preferentially in the vascular tissues for *PtSHR1pro::GUS*, but specifically in the boundary zone between the SAM and leaf primordia for *PtSHR2Bpro:GUS* (Fig. 1B).

**Fig. 1.**
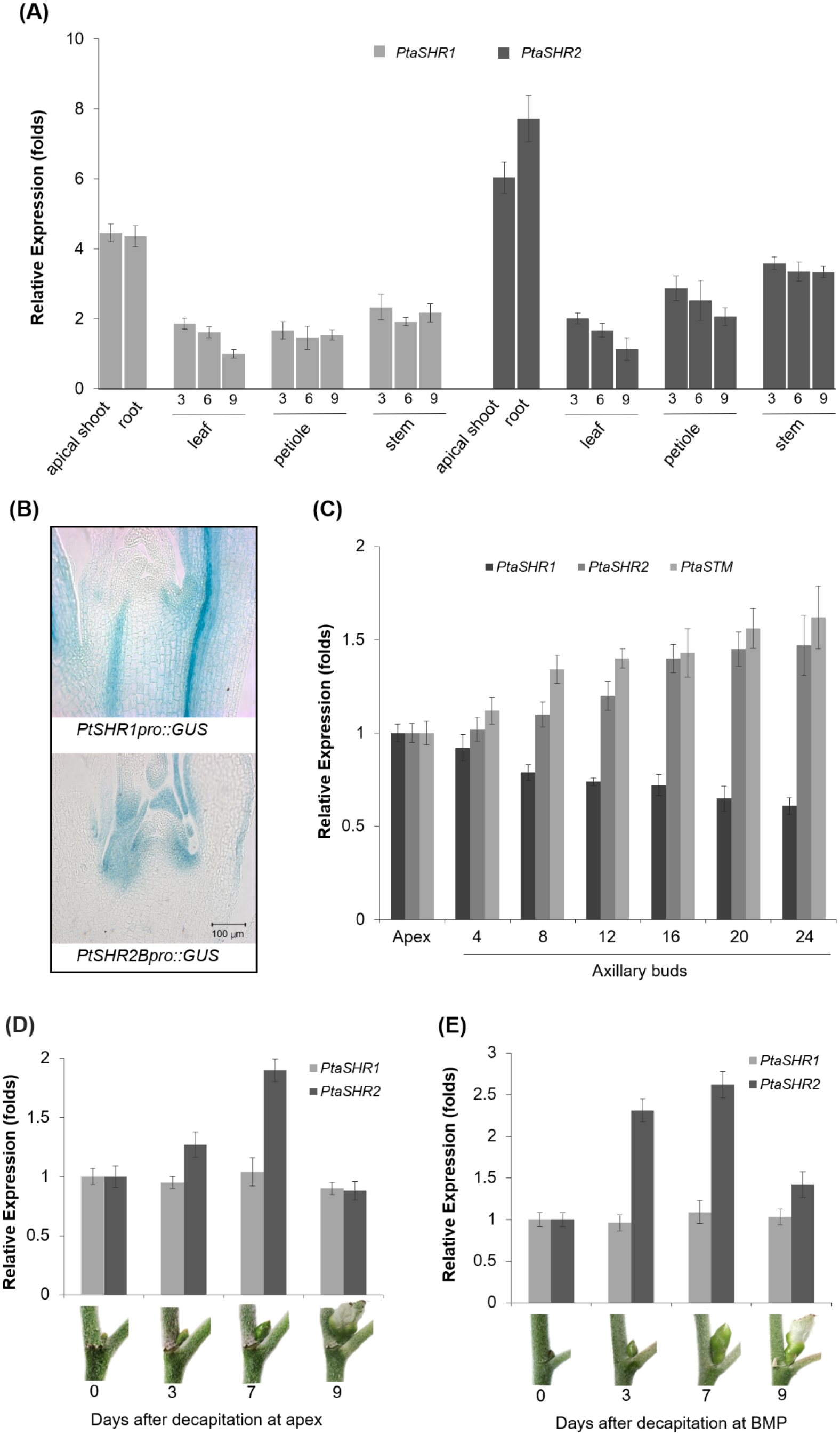
Expression profiles of two *SHR* genes in hybrid poplar (*Populus tremula* × *O. alba* clone INRA 717-1B4). (A) The relative transcription levels of *PtaSHR1* and *PtaSHR2* genes in different tissues of 3-month-old poplar trees by qRT-PCR. Gene expression level of *PtaSHR1* in leaf at the 9^th^ node was set to 1. The experiments were repeated three times, and error bars indicate SD with *n* = 3. (B) GUS-staining assays of two P*taSHR* promoters in poplar shoot apices. Scale bar = 100 μm. (C) Relative expression level of *PtaSHR1* and *PtaSHR2* genes during AXB development. Gene expression level in apex was set to 1. (D) The relative expression of *PtaSHR1* and *PtaSHR2* genes in the nearest AXB at 0, 3, 7, 9 days after decapitation at shoot apex. (E) The relative expression of *PtaSHR1* and *PtaSHR2* genes in the nearest AXB at 0, 3, 7, 9 days after decapitation at the BMP.

To define the spatial and temporal patterns of expression of *PtaSHRs* during AXB development, their expression was analyzed in the proliferating apex and subsequent axillary buds at various nodes. When the expression level of *PtaSHR1* kept a moderate but consistent decrease from the apical shoot down the main stem, *PtaSHR2* revealed increasingly higher expression levels towards the BMP and remained virtually the same since after (Fig. 1C). Such an increasing expression in consecutive AXBs resembled that of an early axillary meristem marker, *SHOOT MERISTEM-LESS* (*PtaSTM*), and implied a putative role of *PtaSHR2* during the process of AXB maturation (Fig. 1C). On contrary, the expression level of two robust, universal markers of the SAM activity (Brand *et al*., 2000), *PtaCLV3* and *PtaWUS*, declined rapidly in developing AXBs below the SAM (Supporting Information Fig. S1).

Decapitation is a process that can release the apical dominance and induce the outgrowth of axillary shoots from the proximal positions. We then investigated the induction patterns of *PtaSHR1* and *PtaSHR2* by decapitation. As shown in Fig. 1D, when decapitation was performed at the apex, the compact and small AXB located underneath grew larger and the tightly packed embryonic leaves subsequently expanded. Within one week, leaf tips became visible and then new leaves totally protruded out. During this process, the expression of *PtaSHR1* at the nearest AXB remained largely unchanged, whereas *PtaSHR2* was transcriptionally upregulated during the first week and then declined to the basal level (Fig. 1D). When decapitation was performed at the BMP (the 16^th^ nodal position), *PtaSHR1* remained unresponsive, whereas *PtaSHR2* again displayed an evident induction at earlier days and then declined (Fig. 1D). Taken together, the AXB-specific expression, the upregulation during bud maturation and the induction in outgrowing buds by decapitation were all suggestive of *PtaSHR2* as a regulator of AXB development.

### *PtSHR2B* overexpression leads to axillary bud outburst below the BMP in hybrid aspen

To gain further insight into *SHR2* function, the hybrid poplar clone INRA 717-1B4 was transformed to overexpress *PtSHR2B* under the control of the *CaMV* 35S promoter, which was referred as *PtSHR2*^OE^ trees. Distinct transgenic phenotypes such as curling and clenched leaves, twisted petioles and swollen stems have been observed during the shoot regeneration process, although these phenotypic abnormalities became much less evident after the rooting stage (Fig. 2A). With dramatically increased transcript abundances of *PtSHR2* (Fig. 2B), both *in vitro* cultured plantlets and transgenic trees grown in greenhouses displayed obvious dwarfness (Fig. 2C-D). Most noteworthy, when *PtSHR2B*^OE^ plants were transferred to soil, they developed sylleptic branches about 3 months later coincident with the transition of the shoot meristem into a terminal bud (Fig. 3A-B). By contrast, all axillary buds of wild-type trees at this stage stayed arrested due to the apical dominance exerted by an actively growing SAM (Fig. 3A-B).

**Fig. 2.**
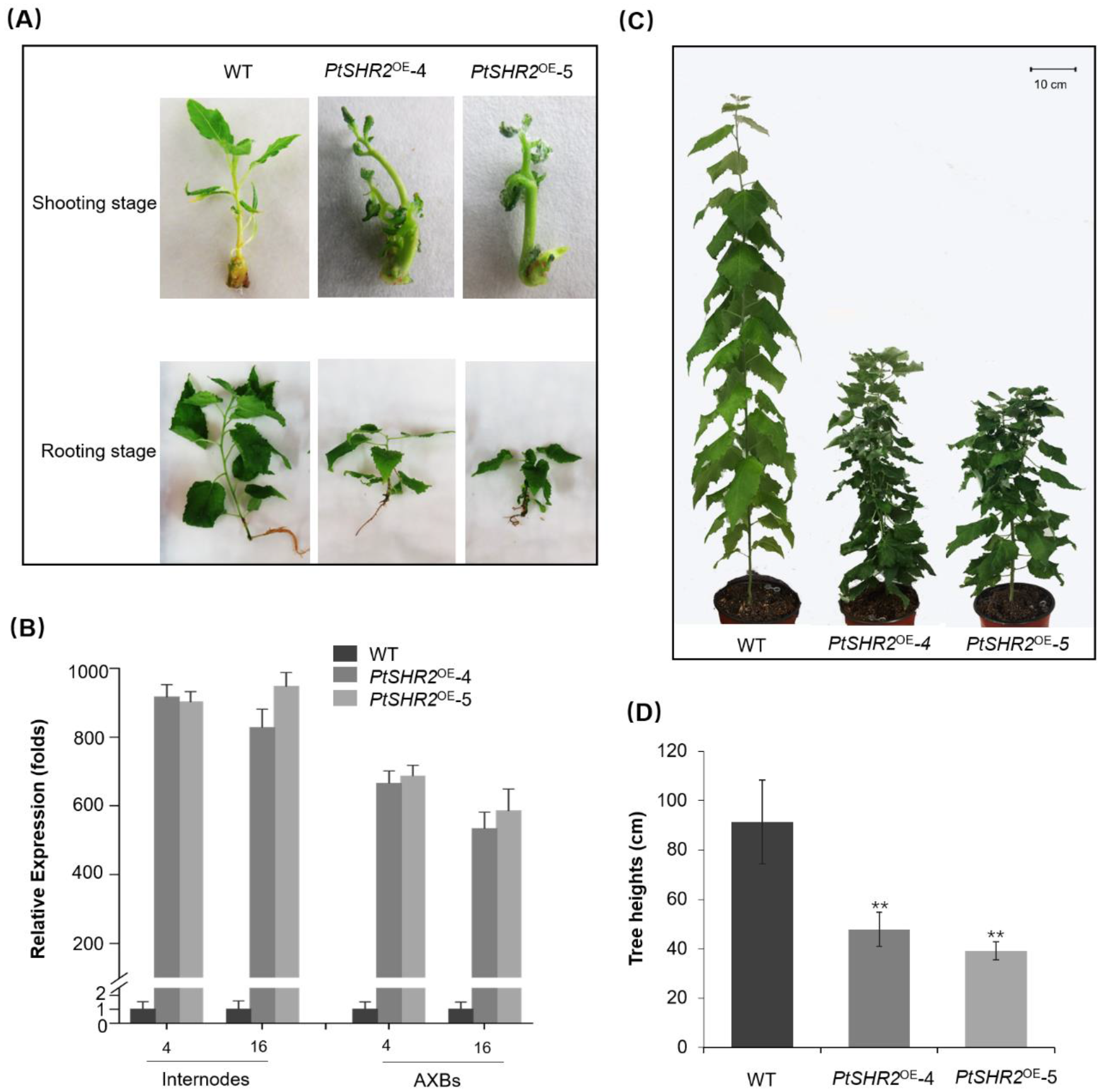
Phenotypic characters of *PtSHR2*^OE^ poplars. (A) Characteristics of *PtSHR2*^OE^ poplar at transformation stages. (B) The relative transcription levels of *PtaSHR2* gene in different tissues analyzed by qRT-PCR. Tissue samples were taken from 2-month-old WT and *PtSHR2*^OE^ trees. The experiments were repeated three times, and error bars indicate SD with *n =* 3. (C) Representative photographs of *PtSHR2*^OE^ transgenic and WT poplars in soil for 4 months. (D) Heights of *PtSHR2*^OE^ transgenic and WT poplars in soil for 4 months. * and ** indicate significant differences between WT and *PtSHR2*^OE^ determined by Student’s t-test at *P* < 0.05 and *P* < 0.01, respectively.

**Fig. 3.**
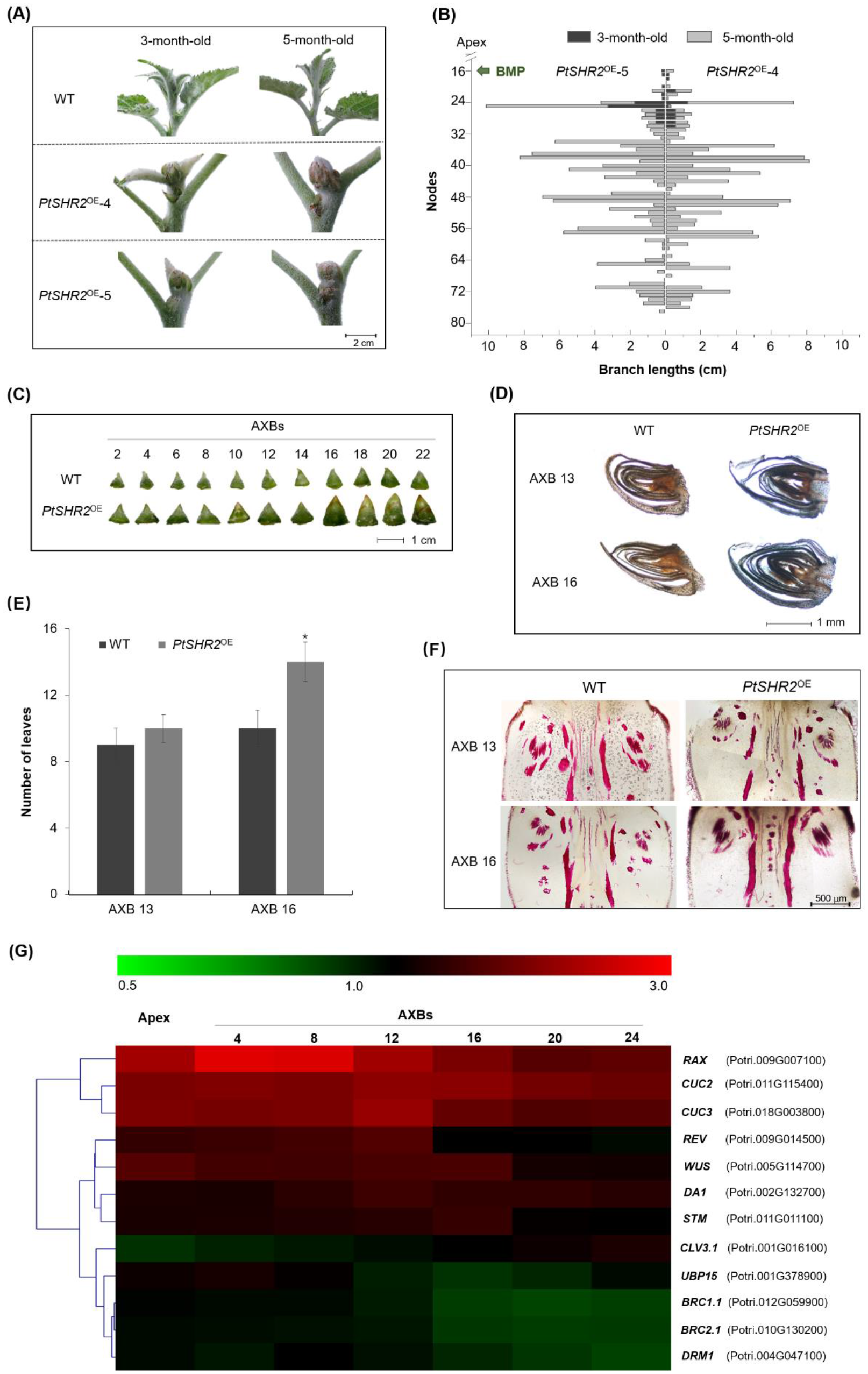
AXBs characters and gene expression analyses of *PtSHR2*^OE^ poplars. (A) Representative images of shoot apices of 3-month-old WT and *PtSHR2*^OE^ poplars. Scale bar = 2 cm. (B) Lateral branch lengths of *PtSHR2*^OE^ poplars at 3 months and 5 months. (C) Sizes of AXBs at subsequent nodes of 3-month-old WT and *PtSHR2*^OE^ poplars. Scale bar = 1 cm. (D) Representative images of longitudinal sections of AXB 13 and 16 of 3-month-old WT and *PtSHR2*^OE^ poplars. Scale bar = 1 mm. (E) The number of embryonic leaves at AXB 13 and 16 of 3-month-old WT and *PtSHR2*^OE^ poplars. The experiments were repeated three times, and error bars indicate SD with *n =* 6. * indicate significant differences between WT and *PtSHR2*^OE^ determined by Student’s t-test at *P* < 0.05. (F) The longitudinal sections showing the vascular structure of AXB 13 and 16 stained by phloroglucinol. Scale bar = 500 μm. (G) Relative expression level of genes regulating axillary meristem imitation and bud outgrowth. Tissues were sampled from 3-moth-old WT and *PtSHR2*^OE^ poplars. Gene expression level in WT trees was set to 1. Gene expression was measured by qRT-PCR using the *Actin* gene as the reference and the results were shown as a heat map with color scale at the top of dendrogram representing log2 values and experiments were performed in triplicates.

In proleptic poplar trees, AMs are initiated in an acropetal order and buds nearest to the apex were younger than buds farther from the apex, so that a gradient of developmental stages was found along the nodes. AXB outburst in *PtSHR2*^OE^ trees unambiguously occurred to nodes below the BMP (Fig. 3A-3B), in concert with the acropetal activation of basal buds often observed in nature. The size of transgenic AXBs was consistently larger than those of WT trees at the same nodal location, particularly at and below BMP (Fig. 3C). In longitudinal sections, the number of embryonic leaves in transgenic AXBs had increased from 9 to 10 at node 13 and from 10 to 14 at node 16 (Fig. 3D-E). A more fully developed vascular structure connecting to the primary stem, which might facilitate the auxin export from AXBs, was also detectable in *PtSHR2*^OE^ AXBs (Fig. 3F). Thus, overexpression of *PtSHR2* seemed to have accelerated the development and maturation of AXBs prior to their escape of paradormancy under a weakened apical dominance.

In woody perennials, very little is known about the molecular processes that control branching, contrary to the intensive investigation in herbaceous Arabidopsis (Katyayini *et al*., 2019). We then searched the poplar genome for homologs of branching regulators in *Arabidopsis* and analyzed their expression changes in transgenic apices and AXBs. First, *PtaSHR2*^OE^ apical shoots revealed lower and higher transcript abundances of *PtaCLV3* and *PtaWUS*, respectively, indicating an interfered apical meristem cell maintenance (Fig. 3G). Second, the expression levels of poplar homologs of transcription factors including *STM*, *REVOLUTA* (*REV*), *REGULATOR OF AXILLARY MERISTEMS* (*RAX*), *CUP-SHAPED COTYLEDON* (*CUC2* and *3*) and *DA1* encoding a ubiquitin-activated peptidase (Greb *et al*., 2003; Li *et al*., 2020; Raman *et al*., 2008; Tian *et al*., 2014; Wang and Jiao, 2018; Yang *et al*., 2012) were all distinctly regulated in *PtSHR2*^OE^ apices and axillary buds (Fig. 3G). *PtaCUC2*/3 and *PtaRAX* genes were highly upregulated at apex and in AXBs, particularly those upper the BMP, while the expression of *PtaREV*, *PtaWUS*, *PtaSTM* and *PtaDA1* were increasingly elevated and reached their maximum in mature AXBs. All these genes have been demonstrated to be essential at early stages of AM initiation to give rise to axillary buds at the base of leaves. Third, the mRNA abundances of *BRANCHED1* (*PtaBRC1*) and *PtaBRC2* that encode branching repressors (Aguilar-Martínez et *al.*, 2007) and DORMANCY ASSOCIATED PROTEIN 1 (*PtaDRM1*) that encodes an early marker of bud dormancy in multiple species (Kebrom *et al*., 2006; Stafstrom *et al*., 1998; Tatematsu *et al*., 2005) were significantly reduced in transgenic AXBs with the most prominent decrease occurring at the BMP and thereafter gradually recovered (Fig. 3G).

### Hormonal homeostasis and signaling were modulated in *PtSHR2*^OE^ hybrid poplar

Once initiated, the fate of an axillary bud is controlled by internal hormonal signals. Considering the crucial role of auxin in repressing the outgrowth of AXBs, we first measured the endogenous IAA contents in *PtSHR2*^OE^ trees. At transgenic shoot apices, the IAA level was 18% higher than that of wild-type plants (Fig. 4A) without upregulating auxin biosynthetic genes (Fig. S2). In contrast, IAA levels in transgenic AXBs as well as adjacent stem segments were 23% lower than those in wild type trees (Fig. 4A). When the expression levels of genes involved in auxin response and signaling were analyzed, members of the auxin/indole-3-acetic acid (Aux/IAA) coreceptor family including *PtaIAA14* and *PtaIAA16* were highly upregulated in transgenic buds near the BMP, but not in the younger nodes farther up (Fig. 4B). IAA14 is a well-known regulator of auxin-regulated lateral root formation and its gain of function mutation was associated with decreased expression of auxin-inducible genes (Fukaki *et al*., 2002). Similarly, a dominant gain-of-function mutation in *IAA16* confers dramatically reduced auxin responses in a variety of assays (Rinaldi *et al*., 2012). The auxin response gene *auxin-resistant1* (*PtaAXR1*) was found to be downregulated at the BMP, and such a reduced bud auxin response was consistent with that the recessive Arabidopsis *axr1* mutant has weaker apical dominance and shows an enhanced shoot-branching potential at maturity (Lincoln *et al*., 1990; Stirnberg *et al*., 1999). In addition, two AXB-expressing auxin-inducible GRETCHEN HAGEN 3 (*GH3*) genes, *PtaGH3.1* and *PtaGH3.6* (Fig. 4B), were also downregulated in transgenic buds, which was consistent with the decreased endogenous auxin levels.

**Fig. 4.**
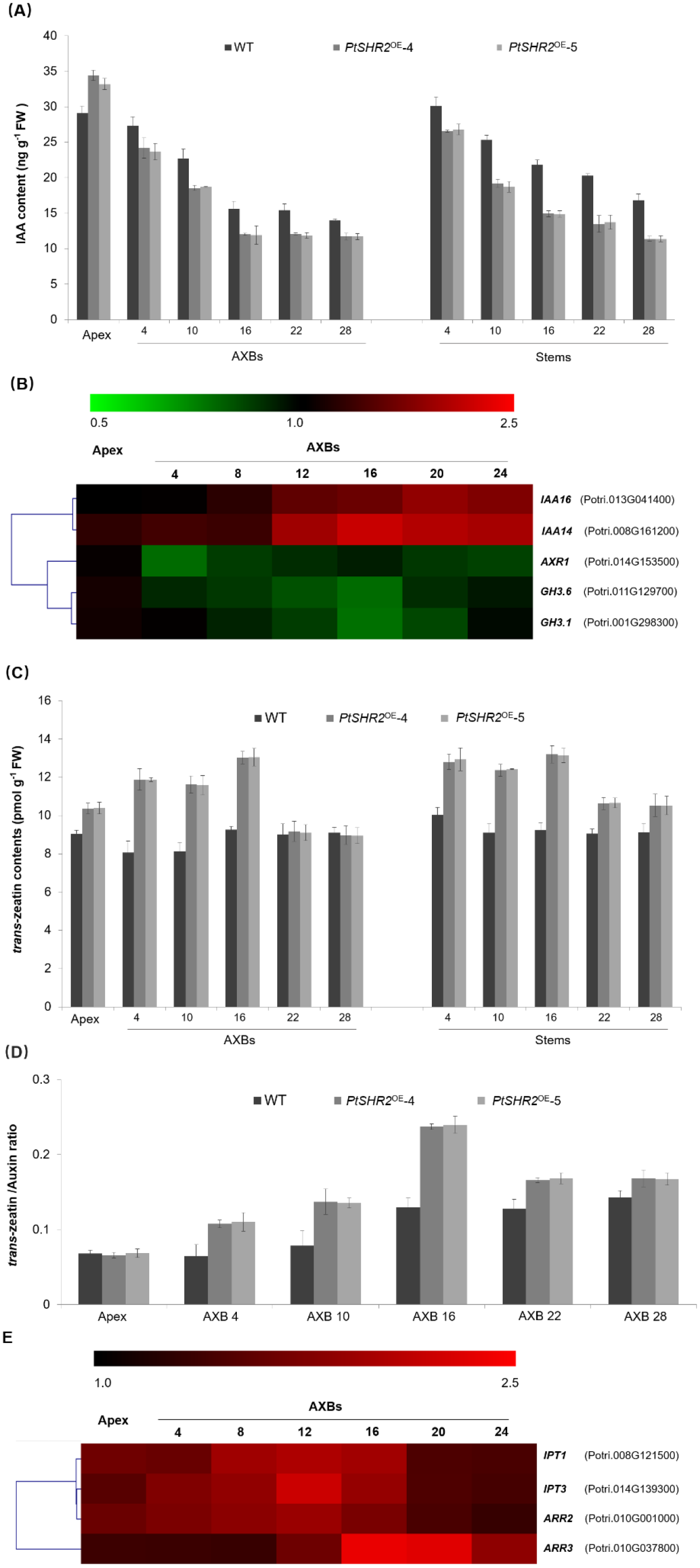
Comparison of hormone levels and expression levels of genes involved un signaling transduction in WT and *PtSHR2*^OE^ hybrid poplars. (A) The endogenous IAA contents in apex, axillary buds, and stems of 3-month-old WT and *PtSHR2*^OE^ poplars. The experiments were repeated three times, and error bars indicate SD with *n =* 3. (B) Gene expression level of auxin-responsive genes in poplar AXBs by qRT-PCR analyses. Tissues were sampled from 3-month-old WT and *PtSHR2*^OE^ poplars. Gene expression level in WT trees was set to 1. The results were shown as a heat map with color scale at the top of dendrogram representing log2 values and experiments were performed in triplicates. (C) The *trans*-zeatin contents in apex, axillary buds, and stems of 3-month-old WT and *PtSHR2*^OE^ poplars. The experiments were repeated three times, and error bars indicate SD with *n =* 3. (D) Ratio of *trans*-zeatin to auxin. The experiments were repeated three times, and error bars indicate SD with *n =* 3. (E) Gene expression level of cytokinin biosynthetic genes in AXBs by qRT-PCR analyses. Gene expression level in WT trees was set to 1. The results were shown as a heat map with color scale at the top of dendrogram representing log2 values and experiments were performed in triplicates.

We then measured the amount of *trans*-zeatin in *PtSHR2*^OE^ hybrid poplars and found that compared with the control, *trans*-zeatin in AXBs was significantly increased and reached the maximum level at the BMP (Fig. 4C), exactly where lateral branches began to emerge out. The ratio of cytokinin to auxin has been regarded as a strong determinant of the degree of lateral shoot outgrowth (Chatfield *et al*., 2000) and it increased with the maturity of axillary buds in both wild-type and transgenic trees. However, except for apex, this ratio in each transgenic AXB was significantly higher than that in its wild-type counterpart (Fig. 4D). Cytokinin biosynthetic genes including *ISOPENTENYL TRANSFERASE1* (*PtaIPT1*) and *PtaIPT3*, as well as type-A Arabidopsis response regulators (*PtaARR3*) and type-B ARR (*PtaARR2*), were all transcriptionally upregulated in transgenic apex and AXBs (Fig. 4E). Type-A ARRs are prime cytokinin response genes with inducibility by cytokinin and *MsARR3/5* have been shown to be required for CK-mediated bud outgrowth in apple trees (Tan *et al*., 2019). Type-B ARRs are transcription factors acting as mediators of cytokinin responses and Arabidopsis ARR2 has been previously shown to directly bind and activate the expression of *WUS* to define the stem cell niche during axillary meristem formation (Meng *et al*., 2017; Wang *et al*., 2017).

### Effects of auxin manipulation on branching of *SHR2*^OE^ lines

Defoliation is known to deplete auxin within stem and thus induces bud outgrowth. With a reduced auxin level in stem, we tested whether *PtSHR2*^OE^ hybrid poplar retained the susceptibility to defoliation. after 30 days, post-decapitation bud outgrowth in control and transgenic plants reached 11.8 cm and 20.9 cm in total, respectively (Fig. 5A and 5B). Compared to nondefoliated transgenic trees, defoliation did not significantly affect the kinetics of bud activation or activate significantly more buds (Fig. S3), indicating that defoliation did not enhance the branching pattern of *PtSHR2*^OE^ transgenic trees. Decapitation is another classical assay to study bud activation after release from apical dominance (Tatematsu *et al*., 2005), we then tested the responsiveness of transgenic trees to decapitation at three sites (below apex, at BMP and at the basal node) and observed more expeditious bud enlargement and leaf outgrowth in *PtSHR2*^OE^ lines than in wild type trees, and the closer to the base of the stem, the more pronounced the difference (Fig. 5C-5D).

**Fig. 5.**
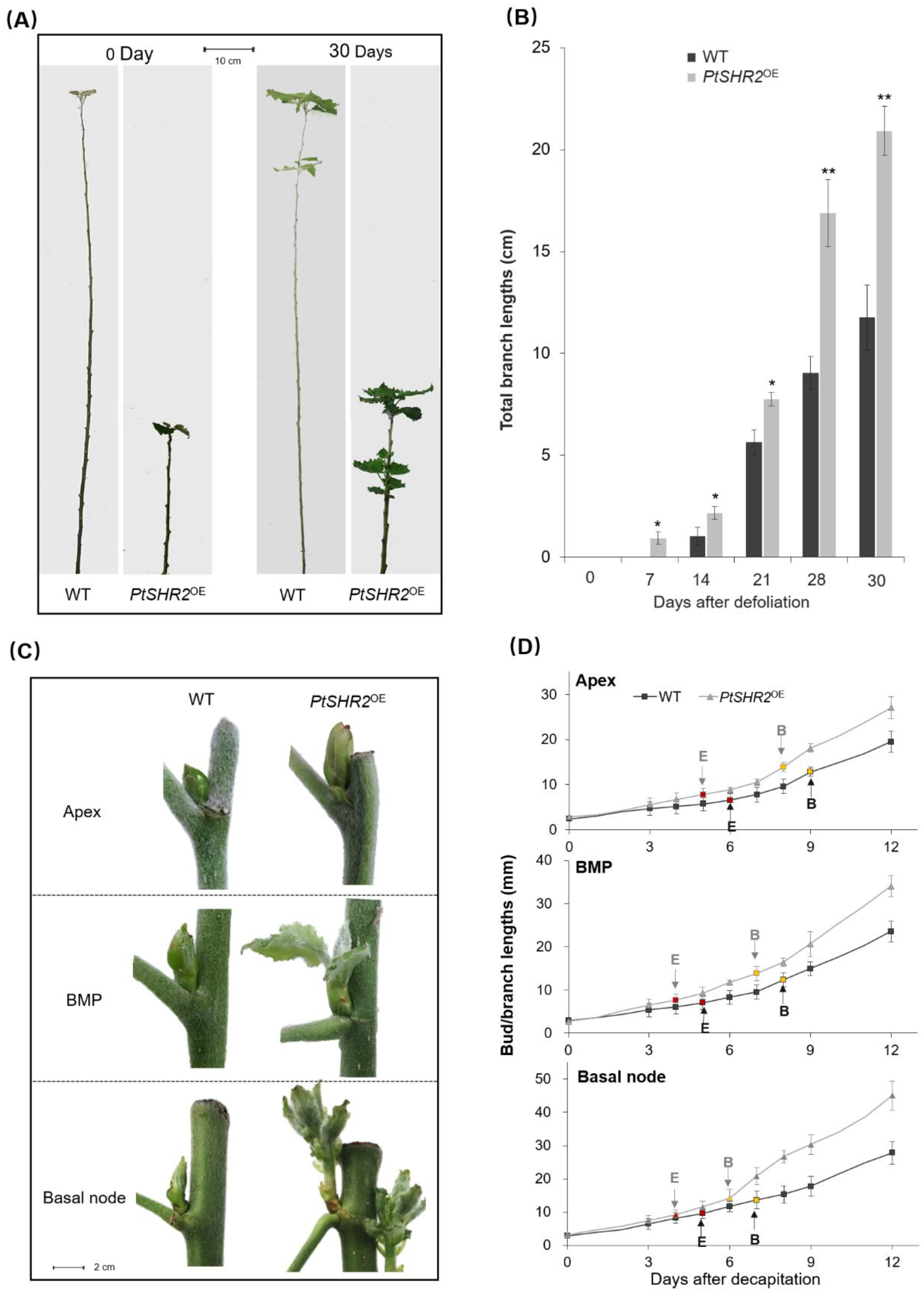
Defoliation and decapitation responses of AXBs in 3-month-old wild-type and transgenic hybrid poplars. (A) Representative images showing the responsiveness of WT and *PtSHR2*^OE^ poplars after defoliation at day 0 and 30. Scale bar = 10 cm. (B) Total branch lengths of WT and *PtSHR2*^OE^ poplars after defoliation after a 30-day period. Error bars indicate SD with *n =* 6. * and ** indicate significant differences between WT and *PtSHR2*^OE^ determined by Student’s t-test at *P* < 0.05 and *P* < 0.01. (C) Representative images showing the responsiveness of WT and *PtSHR2*^OE^ poplars in the nearest bud after decapitation at three sites (apex, the BMP and the basal node) at day 8. Scale bar = 2 cm. (D) The total bud and branch lengths of the nearest AXB after decapitation at day 0, 3, 6, 9 and 12. The arrow ‘E’ marks the day when AXBs significantly enlarged (E), and the arrow ‘B’ marks the AXB burst. Error bars indicate SD with *n =* 6.

NAA, at concentrations as high as 5 mg g^−1^ in lanolin, has been demonstrated to be able to increase the endogenous IAA content to above that of intact control plants (Morris *et al*., 2005) and lower concentrations have been reported to have less effective at inhibiting bud outgrowth (Beveridge *et al*., 2000). When NAA was applied to the shoot apex following decapitation, it effectively diminished the outgrowth of the first and second nearest buds in transgenic trees without disturbing the emergence of AXBs below the BMP (Fig. 6A-6B, S4). Notably, the inhibitory effect of NAA on transgenic AXB was even stronger than on WT buds. In the absence of NAA, decapitation at apex induced bud/shoot outgrowth in *PtSHR2*^OE^ transgenic trees as 135% as that of WT at the 1^st^ next node, while with NAA, the relative bud enlargement decreased to only 50% of WT elongation, indicating a higher auxin sensitivity of transgenic AXBs than wild-type AXBs. Such enhanced responsiveness to NAA was also observed below the BMP (Fig. 6C-D). Taken together, *PtSHR2*^OE^ AXBs maintained a normal even higher sensitivity towards exogenous auxin, in which case the major factor responsible for the activated AXBs seemed to be an inefficient auxin basipetal transport in transgenic stems, which in turn\ caused an auxin accumulation at the transgenic apex.

**Fig. 6.**
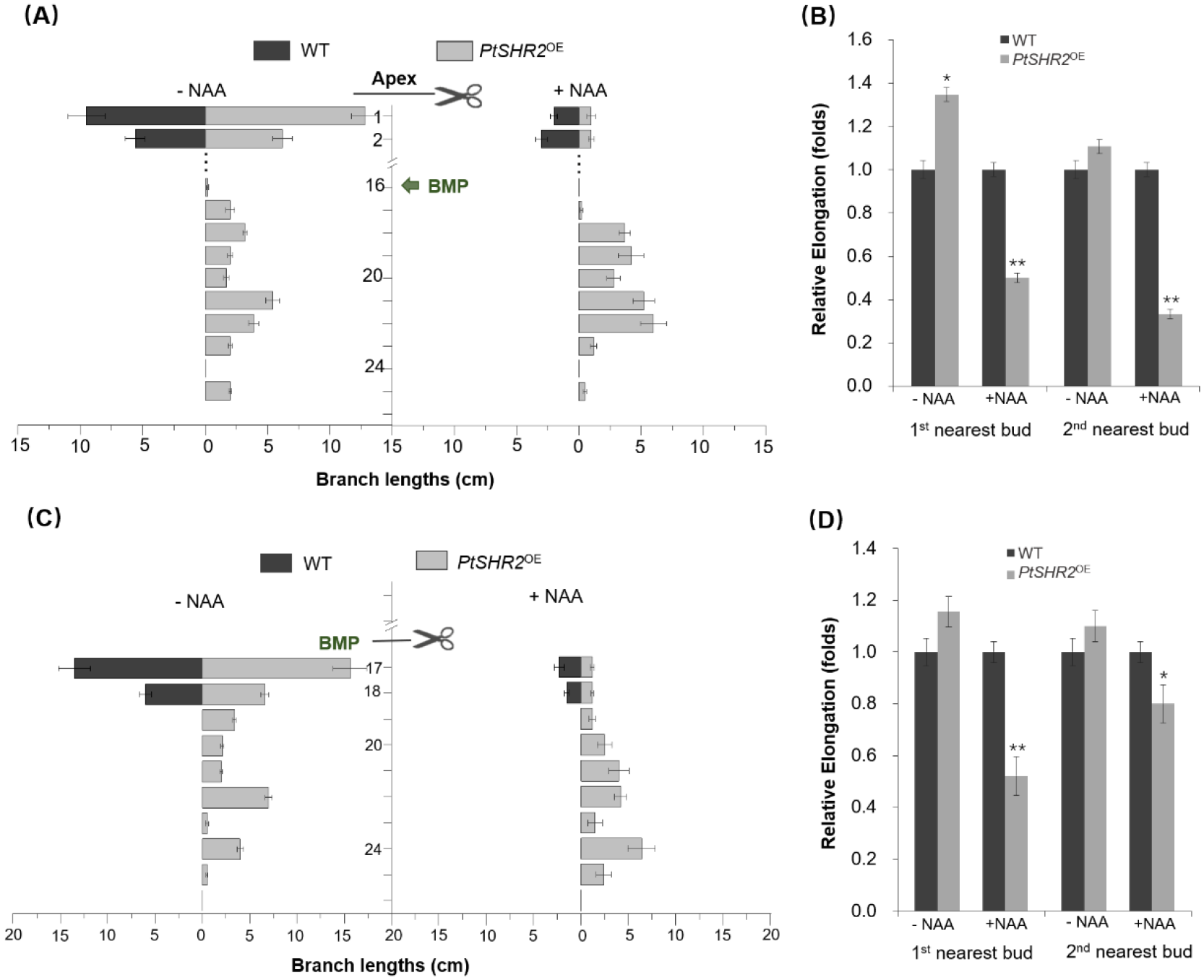
Effects of NAA on *PtSHR2*^OE^-enhanced axillary bud outgrowth. (A) Branch lengths of 3-month-old WT and *PtSHR2*^OE^ poplars supplemented with 6 mg g^-1^ NAA at the decapitated sites at shoot apex after 30 days. Error bars indicate SD with *n =* 6. (B) The relative elongation of bud and branch outgrowth in the 1^st^ and 2^nd^ nearest buds of 3-month-old WT and *PtSHR2*^OE^ poplars supplemented with or without 6 mg g^-1^ NAA in lanolin on the decapitated shoot apices for 30 days. Error bars indicate SD with *n =* 6. * and ** indicate significant differences between WT and *PtSHR2*^OE^ determined by Student’s t-test at *P* < 0.05 and *P* < 0.01. (C) Branch lengths of 3-month-old WT and *PtSHR2*^OE^ poplars supplemented with 6 mg g^-1^ NAA at the decapitated sites at BMP after 30 days. Error bars indicate SD with *n =* 6. (D) The relative bud and branch outgrowth in the 1^st^ and 2^nd^ nearest buds of 3-month-old WT and *PtSHR2*^OE^ poplars after decapitation at the BMP and the following supplement of 6 mg g^-1^ NAA in lanolin for 30 days. Error bars indicate SD with *n =* 6. * and ** indicate significant differences between WT and *PtSHR2*^OE^ determined by Student’s t-test at *P* < 0.05 and *P* < 0.01.

### Effects of PAT and cytokinin biosynthesis inhibitors on *SHR2*^OE^ bud burst

Cytokinin has been proposed to regulate shoot branching by exporting auxin from buds through auxin canalization (Bennett *et al*., 2016). In Arabidopsis, PIN1 is an important component of the classical polar auxin transport (PAT) responsible for the bulk polar movement of auxin through the stem (Petrasek *et al*., 2006), while PIN3, PIN4, and PIN7 are major contributors to a less polar route, termed connective auxin transport (CAT), which connects the surrounding tissues including axillary buds with PAT and regulates shoot branching (Bennett *et al*., 2016). In transgenic AXBs, the transcript levels of *PtaPIN1*, *PtaPIN3*, *PtaPIN4* and *PtaPIN7* were all drastically upregulated (Fig. 7A), implicative of a mobilized PAT system in *PtSHR2*^OE^ buds.

**Fig. 7.**
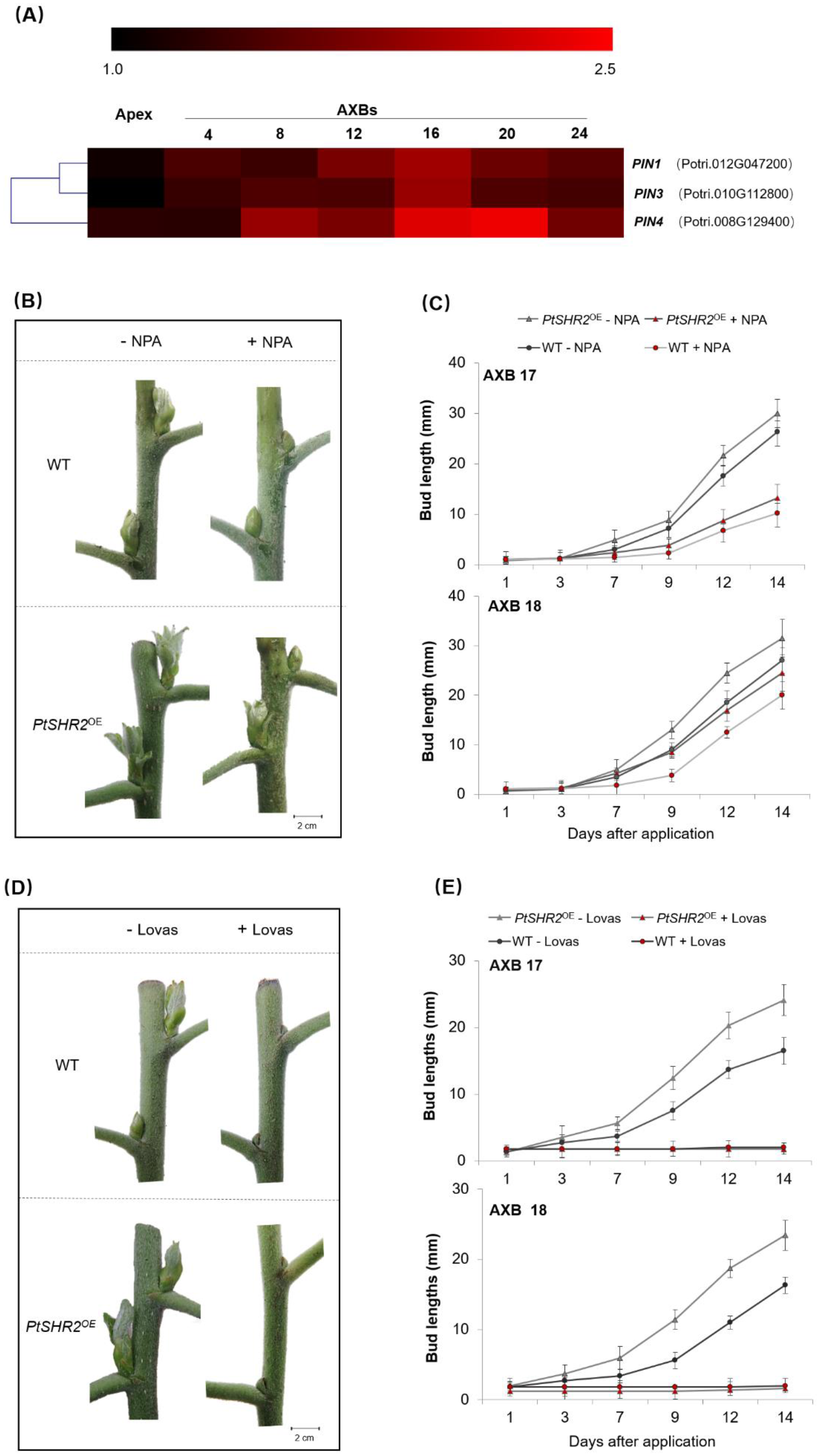
Effects of NPA and Lovastatin on *PtSHR2*^OE^-enhanced axillary bud outgrowth. (A) Gene expression levels of *PIN* genes by qRT-PCR analyses. Tissues were sampled from 3-month-old WT and *PtSHR2*^OE^ AXBs. The experiments were performed in triplicates and error bars indicate SD with *n =* 3. (B) Representative images showing the responsiveness of 3- month-old WT and *PtSHR2*^OE^ AXBs after application of 30 μM NPA onto the decapitated stumps at day 7. Scale bar = 2 cm. (C) The outgrowth of AXB17 and AXB18 measured at day 14 after 30 μM NPA application. Error bars indicate SD with *n =* 6. (D) Representative images showing the responsiveness of 3- month-old WT and *PtSHR2*^OE^ AXBs after application of 5 mM lovastatin onto the decapitated stumps at day 7. Scale bar = 2 cm. (E) The outgrowth of AXB17 and AXB18 measured at day 14 after 5 mM lovastatin application. Error bars indicate SD with *n =* 6.

We then assessed the possible involvement of PAT in hyperbranching of *PtSHR2*^OE^ by applying 30 μM 1-N-naphtylphtalamic acid (NPA) onto decapitated stumps and investigated the changes in the decapitation-induced activation of proximal buds. In a 12-day time frame, NPA significantly delayed, albeit not completely stop, the first next AXB outgrowth in both wild-type and transgenic trees; and had much less inhibition on the second downward AXB (Fig. 7B-C). Notably, although the NPA-fed wild-type AXBs showed slightly delayed bud initiation, the rate of shoot elongation afterwards were not significantly different from that of transgenic shoots, implying that NPA-feeding might only affect the bud mobilization. In contrast, when Lovastatin, an inhibitor of cytokinin biosynthesis, was applied to the decapitated stumps in the same manner as NPA, it inhibited the high-branching phenotype as completely as NAA did (Fig. 7D-E). We also smeared NPA or Lovastatin directly to transgenic AXBs, in which case NPA turned out to be ineffective but Lovastatin again caused a total inhibition (Fig. S5). Together, these results indicated that the activation and emergence of transgenic AXBs indispensably requires the biosynthesis of cytokinin inside the bud and also need the auxin transport capacity undertaken by multiple PIN proteins, in accordance with recent results in Arabidopsis showing that cytokinin regulation of PIN3/4/7 proteins is not the only mechanism by which cytokinin promotes shoot branching (Waldie and Leyser, 2018).

## Discussion

Shoot branching is caused by the activity of axillary meristems (AMs), which are initiated post-embryonically in leaf axil and develop into small axillary buds (AXBs) that in most cases stop growing and enter dormancy due to apical dominance (Domagalska and Leyser, 2011; Martín-Fontecha *et al*., 2018). Different from Arabidopsis, AXMs in most trees arise at a very early stage in the axils of emerging leaves and produce complex AXBs with an enclosed embryo shoots (ES) and sturdy bud scales. In poplars, spontaneous bud burst is rare under normal growth conditions and an ES only give rise to proleptic branches after a winter dormancy (Aguilar-Martínez *et al*., 2007). In this work, the overexpression of *PtSHR2* in transgenic hybrid poplar resulted in a bushier phenotype by affecting the decision of mature axillary buds below the BMP to grow out without delay with a tightly packed rosette of developed embryonic leaves.

The function of SHR involved in AXB formation and outgrowth has not been elucidated in literature, but some pieces of expression evidence in this work proposed the role of poplar *SHR2B* gene during these processes, such as its distinct expression in the region between leaf primordia and the SAM where later form leaf axils, its spatiotemporal regulation during AXB maturation along the stem and its inducible expression in activated AXBs after decapitation (Fig. 1A-E). Moreover, *PtSHR2*^OE^ trees displayed a high branching phenotype at about 3 months of age, and the emergence of side branches was always accompanied by a gradually terminated apex, pointing to an identity exchange between the apical meristem and the axillary buds (Fig. 3A-B). This observation was consistent with previous finding that removal of apical dominance in hybrid aspen could shift the gene expression of AXBs toward that of apex, while developing terminal buds (TBs) adopted the expression pattern of paradormant AXBs (Rinne *et al*., 2015). Intriguingly, the occurrence of such identity transition was not random; instead, the sustained bud outgrowth of *PtSHR2*^OE^ trees was confined to a subset of AXBs located below the BMP, suggesting that buds’ maturation status was a major phenotypic determinant. Indeed, transgenic AXBs were evidently larger and included more embryonic leaves and more developed vascular tissues when compared with wild-type buds (Fig. 3C-E), suggesting their accelerated axillary meristem initiation and bud maturation.

The coordinated and partially redundant actions of a series of transcription factors have been implicated in the formation of the secondary shoot meristem from a small group of cells in the leaf axil (Schmitz and Theres, 2005). For instance, STM (Greb *et al*., 2003; Shi *et al*., 2016) and REV (Lee *et al*., 2009; Otsuga *et al*., 2001) are required for AM initiation and maintenance. NAC transcriptional factors CUC2 and CUC3 participate in the boundary formation and AM initiation during postembryonic development (Aida *et al*., 1997; Hibara *et al*., 2006; Raman *et al*., 2008). Synergistic interaction between DA1 and a MYB transcription factor, RAX1, regulates the AM initiation during vegetative and reproductive development (Keller *et al*., 2006; Muller *et al*., 2006). All these transcriptional factors were significantly upregulated in their mRNA abundances in transgenic apex and AXBs, although they followed different trends in successive axillary buds (Fig. 3G). The joint actions of all these signals may constitute a coordinated gene regulatory network through different pathways to enhance the initiation and establishment of AXM in the center of *PtSHR2B*^OE^ leaf axil. Meanwhile, the well-known integrator of signals controlling bud outgrowth, BRANCHED1 (BRC1) and BRC2, which act exclusively in axillary buds as a conserved branching suppressor (Aguilar-Martínez *et al*., 2007) were down-regulated in transcription, together with another branching repressor, *DRM1* (Stafstrom *et al*., 1998; Tatematsu *et al*., 2005). Notably, upregulation of transcription factors that promote AM initiation occurred primarily in developing AXBs, while down-regulation of transcription factors that inhibit branching was more pronounced in mature AXBs below the BMP, consistent with their respective range of action. Together, the characteristic and differential transcriptional regulations of these key branching regulators paved the way for earlier AM initiation and faster development of *PtSHR2*^OE^ buds.

Once an axillary bud has established, it often arrests as a dormant bud. Auxin, produced in the shoot apex and actively transported basipetally through a specific polar auxin transport (PAT) stream, plays a central role in this process without entering the bud. In contrast to its weak apical dominance, the auxin level at the transgenic shoot apex was not lower but higher than that of wild-type trees, without upregulating the corresponding IAA biosynthetic genes (Fig. 4A, S2). However, in transgenic AXBs as well as their adjacent stem segments, the level of auxin was significantly lower than in control tissues (Fig. 4A), consistent with that constitutive repression of auxin in leaf axils enhances the side shoot formation (Wang *et al*., 2014) and plants with reduced auxin signaling/level display increased branching levels(Booker *et al*., 2003; Romano *et al*., 1991). Auxin response factor 1 (AXR1) is an integral component of the auxin signaling pathway and *axr1-12* mutant promoted bud outgrowth as a result of the loss of auxin sensitivity (Stirnberg *et al*., 1999). In transgenic trees, *PtaAXR* expression was enhanced at apex but decreased at the BMP (Fig. 4B), implicative of the differentially regulated auxin signaling activities at these two locations.

Consistent with the evidence that auxin in the main stem reduces the synthesis of cytokinin and consequently inhibits bud activity (Tanaka *et al*., 2006), an increased cytokinin level was found in transgenic AXBs, culminating in a 130% level of WT at the BMP (Fig. 4C). Thus, it is plausible that due to its inefficient basipetal transport, the auxin in the primary stem of *PtaSHR2*^OE^ trees was not saturated enough to inhibit bud growth by downregulating cytokinin biosynthesis. Elevated cytokinin levels has been previously identified in defoliated tobacco plants (Geuns *et al*., 2001) and in decapitated pea plants (Chabikwa *et al*., 2019). Together, the decreased auxin level and the increased cytokinin level within the transgenic AXBs resulted in a much higher cytokinin/IAA ratio than wild-type AXBs (Fig. 4D), which has been associated with the outgrowth of axillary buds in tobacco (Geuns *et al*., 2001) and two differently branching *Chrysanthemum morifolium* genotypes (Dierck *et al*., 2016). Cytokinin signaling has been shown to activate the *de novo* expression of *WUS* in the leaf axil and thus establish AM through the direct regulation of type-B Arabidopsis response regulators (ARRs) (Wang *et al*., 2017). In this work, the expression level of *PtaWUS* decreased at the top of the transgenic tree, but it gradually increased in subsequent axillary buds (Fig. 4B), which was well correlated with the increase of cytokinin content in these buds (Fig. 4C). Cytokinin can also establish a positive feedback loop with *STM* expression during AM initiation (Rupp et al., 1999) and negatively regulates *BRC1* expression as an early response to the bud release from apical dominance (Braun *et al*., 2012; Minakuchi *et al*., 2010). In summary, compared with wild-type buds, the much higher ratio of cytokinin to IAA in *PtaSHR2*^OE^ AXBs might be the main factor affecting the degree of lateral branch growth as reported previously (Chatfield *et al*., 2000).

Next, we tried to perturb the high branching phenotype of *PtSHR2*^OE^ trees by manipulating its internal hormone homeostasis. First, when the auxin content in the primary stem was further reduced by defoliation, the further promotion of axillary bud outgrowth in *PtSHR2*^OE^ trees was negligible (Fig. 5B). However, transgenic AXBs responded more rapidly to decapitation than wild-type AXBs, as manifested by their earlier outburst and longer branches and the further the decapitation site from the shoot top, the more pronounced the difference (Fig. 5C-D), and such short preparation time could be underlined by the more precocious embryotic shoot development within the transgenic AXBs (Fig. 3C-E).

In terms of response to exogenously applied NAA, the decapitated stumps of *PtSHR2*^OE^ trees not only retained but demonstrated a significantly higher sensitivity towards NAA than wild-type trees (Fig. 6A-D), implying that through decapitation and NAA reapplication, the impaired top-down transport of auxin in transgenic stem could be bypassed through removing the transportation distance from the auxin stock to the sink and thus restored the apical dominance and inhibited the bud outgrowth. The polar auxin transport (PAT) system dictated by IAA efflux carriers, particularly PIN-FORMED1 (PIN1), is suggested to be responsible for the unidirectional flow of auxin through the stem (Friml *et al*., 2003) and so far, there is no clear correlation between the PAT in the stem and shoot branching (Brewer *et al*., 2015). In addition, the local transport of auxin between the PAT stream and surrounding tissues including axillary meristems, termed as connective auxin transport (CAT), is mediated by PIN3, PIN4, and PIN7 proteins. In Arabidopsis, CAT is suggested to not only convey the information about active apex to each AXB on the stem, but also promote auxin transport connectivity by loading auxin from the bud into the PAT in auxin canalization, thus playing an important role in shaping shoot system architecture (Bennett *et al*., 2016). Arabidopsis mutants impaired in CAT display decreased branching (Bennett *et al*., 2016) and PIN3, PIN4, and PIN7 have been reported to regulate shoot branching in a BRC1-independent manner in Arabidopsis (van Rongen *et al*., 2019) or in a BRC1-dependent manner in cucumber (Shen *et al*., 2019). In this work, the transcript abundances of poplar homologs of *PIN1, 3, 4* and *7* were all dramatically increased in AXBs, with the maximum identified at the BMP. In pea, inhibition of auxin export specifically from the bud reduced bud outgrowth (Chabikwa *et al*., 2019). In this work, although NPA had no effect on bud outgrowth when directly smeared onto axillary buds of intact transgenic trees, NPA applied to the decapitated stumps could retard the branch outgrowth of the nearest bud to a moderate degree, indicating auxin export from buds was involved in the bud mobilization and the sustained outgrowth of axillary shoots (Balla *et al*., 2002). Auxin export from the axillary bud is regarded as not essential to trigger initiation of bud growth but is important for sustained bud outgrowth (Barbier *et al*., 2019; Chabikwa *et al*., 2019), in accordance with that after NPA treatment, the elongation of wild-type and transgenic lateral branches was almost the same (Fig. 7B-C). Cytokinin has been reported to act independently to regulate the growth of axillary buds rather than as a second messenger for auxin (Chatfield *et al*., 2000), although in Arabidopsis, decapitation still releases the buds of cytokinin deficient or resistant mutants from apical dominance (Müller *et al*., 2015). In contrast, CK inhibitor, Lovastatin, effectively suppressed axillary bud outgrowth induced by decapitation (Fig. 7D-E). In addition, when NAA suppressed only the proximal axillary buds (Fig. 6C), Lovastatin appiled to the decapiated stump inhibited even more distant axillary buds down the stem (Fig. S6), demonstrating that at least in *PtSHR2*^OE^ trees, the local cytokinin boisynthesis is absolutely required for transgeic AXB activation. Thus, data present here not only affirmed the remarkable conservation of mechanisms regulating branching in herbaceous and woody plants, but also pointed out their evolutionary divergence in controlling the branching process.

In conclusion, *PtSHR2*^OE^ poplar represented a good opportunity to study the process of lateral bud activation in woody plants. With an attenuated auxin-mediated bud inhibition, transgenic AXBs below the BMP developed into branches instead of remaining dormant in the axils of leaves, as manifested by the dynamically regulated expression regime of multiple factors and signaling pathways that influence the initiation of AMs and promote the maturation and outgrowth of axillary bud. With an impaired bulk polar movement of auxin down the main stem, auxin continuously produced at the shoot apex was locally accumulated and the enhanced stem sink strength allowed auxin to be exported from lateral buds through the action of PIN efflux carriers. With a decreased amount of auxin within AXBs, the antagonism between auxin and cytokinin was alleviated, and the inhibited biosynthesis of cytokinin by auxin was released. The resultant increased CK/IAA ratio within the bud is then translated into a local response that changes the bud state and promote shoot branching. Together, our findings provide a striking example of communication between shoot apex and axillary buds in the regulation of shoot branching through auxin-cytokinin interplay and feedback loops and demonstrated the flexibility of stem cell niche reconstruction during organogenesis in woody trees.

## Abbreviations

SHR: Shoot-root
AXBs: Axillary buds
AMs: Axillary meristems
BMP: Bud maturation point
PAT: Polar auxin transport
CAT: Connective auxin transport
GUS: β-glucuronidase
IAA: 3-Indoleacetic acid
NAA: 1-naphthlcetic acid
NPA: N-1-naphthylphthalamic acid

## Supplementary data

Supplementary data are available at *JXB* online.

**Table. S1** Primer sequences used in this study for qRT-PCR analysis.

**Fig. S1** Relative expression level of *PtaWUS* and *PtaCLV3* genes in different tissues of 3-month-old poplar trees by qRT-PCR.

**Fig. S2** Gene expression level of auxin biosynthetic genes in apices and the 4^th^ AXB of 3-month-old WT and *PtSHR2*^OE^ poplars by qRT-PCR analyses.

**Fig. S3** Total branch lengths of intact and defoliated 3-month-old *PtSHR2*^OE^ poplars after 30 days.

**Fig. S4** Representative images showing the NAA application on the decapitated sites at apex and BMP of 3-month-old WT and *PtSHR2*^OE^ poplars.

**Fig. S5** Branch lengths of 3-month-old *PtSHR2*^OE^ poplars supplemented with 30 μM NPA (A) and 5 mM lovastatin (B) directly on transgenic AXBs for 7 and 14 days.

**Fig. S6** Branch lengths of *PtSHR2*^OE^ poplars after decapitation at the BMP and the following supplement of 5 mM lovastatin for 14 days.

## Acknowledgements

This work was supported by National Natural Science Foundation (31870572).

## Author contributions

JW conceived research, designed experiments and wrote the manuscript. MY, HY and SY performed experiments and analyzed data.

## Data Availability Statement

All data supporting the findings of this study are available within the paper and within its supplementary materials published online.

